## Supplementary material for "A light tunable differentiation system for the creation and control of consortia in yeast"

#### Outline

I. Bioreactor platform and flow cytometry data analysis

II. Microscopy platform and microscopy data analysis

III. Differentiation system
   a. Construction and characterization (Figure 2)
   b. Model development, model fitting, and parameter estimation (Figure 2)

IV. Growth arrest upon differentiation system (GAuDi)
   a. Construction and characterization (Figure 4)
   b. Model development, model fitting, and parameter estimation (Figure 4)

V. MPC control

VI. Pattern formation setup and results for diverse patterns (Figure 1)

VII. Construction of the two differentiation programs and quantification of species' prevalence in multi-species consortia (Figure 5)

VIII. Note on precautions undertook during handling and manipulation of the light sensitive strains.

### I. Experimental platform, flow cytometry, and data analysis pipeline

We used our previously described turbidostat platform[1] to continuously culture cells and conduct time course experiments (Figure 2, 3, 4, 5). Briefly, the platform allows us to monitor 16 cultures in parallel with regular OD measurements and maintain them at a target cell density. The OD measurements and dilution data were used to estimate the growth rate. Further, culture vessels are equipped with LEDs and can be stimulated independently with light. Samples from the vessels are collected in a 96-well plate and, with the help of a pipetting robot, loaded into the cytometer. The cytometer acquisition is controlled with the help of click and point software. Figure S1.1a gives a general overview of the experimental platform.

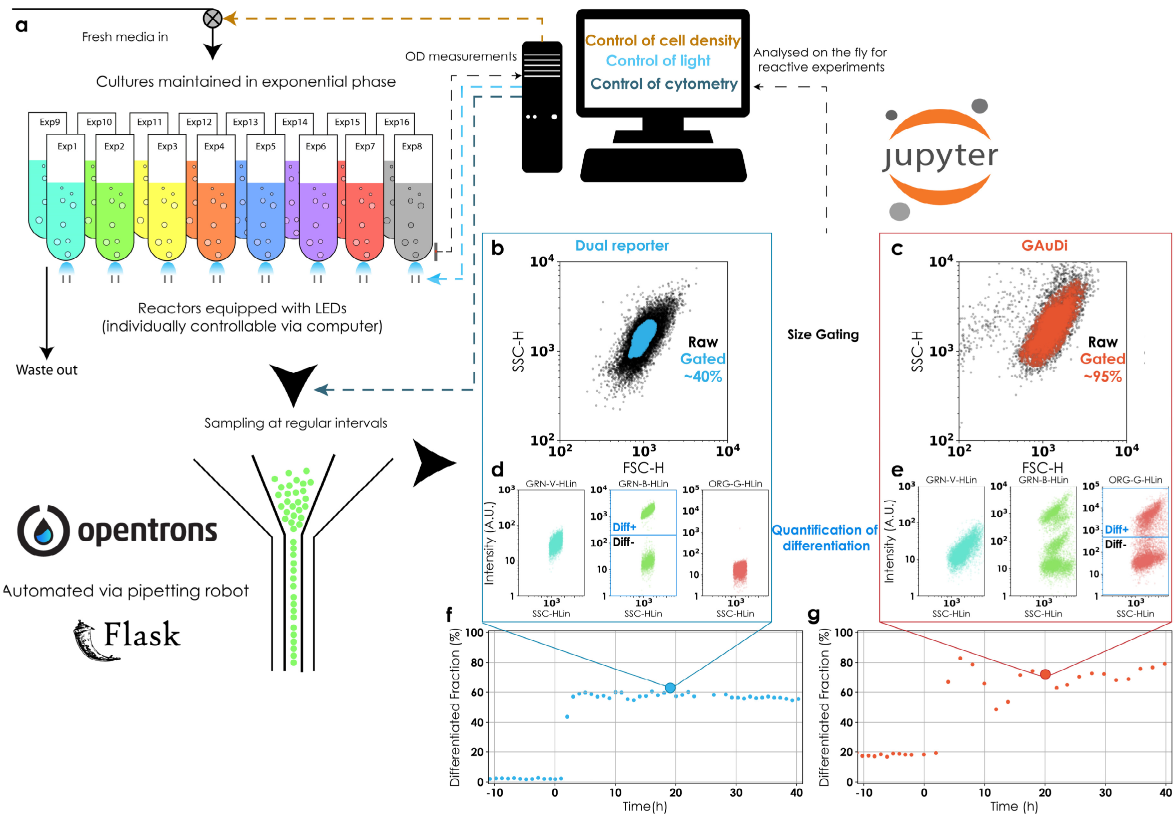


*Figure S1.1 Experimental platform (**a**) and data overview. 16 experiments can be conducted in parallel in reactors with individual OD and LED induction control. The cytometry measurements are automated via a pipetting robot controlled with the Flask app. The raw data from the cytometer is parsed and stored as a csv file or can be analysed online to change experiment conditions such as cell density or induction profiles. All code required for the functioning of the reactors and analysis is developed in Python and implemented using Jupyter*

*notebooks. Representative data from a single timepoint for one experiment each of the original differentiation system (blue box) and GAuDi strain (red box). The FSC vs SSC scatterplot is shown as an indicator of cell size (**b** and **c**). The GRN-B and ORG-G channels are used to detect differentiated cells for the original differentiation system (mNeonGreen fluorescence) and GAuDi (mScarlet-I fluorescence), respectively (blue boxes in **d** and **e**). At this representative timepoint (large circle in **f** and **g**), both exist in two subpopulations.*

All cytometry measurements were made with a Guava EasyCyte BGV 14HT benchtop flow cytometer. Settings and gains were kept constant for all the experiments.

5000 events were recorded for each sample unless specified differently. No compensation was used during acquisition. Dilutions were made with a pipetting robot so that the cell density was kept between 200 (to have 5000 events in the acquisition window) and 600 cells/µl (to ensure >90% singlets). Size gating and doublet removal were done using kernel density based methods. Singlets were selected based on deviation from linearity in Forward Scatter Height (FSC-H) vs. Forward Scatter Area (FSC-A). Cells were scored and a threshold was defined above which cells were classified as doublets and removed from analysis. For size gating, 2D kernel density estimates were obtained using SciPy gaussian kde package on Forward Scatter (FSC-H) vs. Side Scatter (SSC-H) and regions of density lower than a threshold were removed (Figure S1.2). The two thresholds were kept constant for all measurements except those made with the GAuDi strain. In the latter case, thresholds were increased to include the entire population. This leniency was warranted because of considerable changes in the side scatter in growth arrested cells (compare Figure S1.1b & Figure S1.1c).

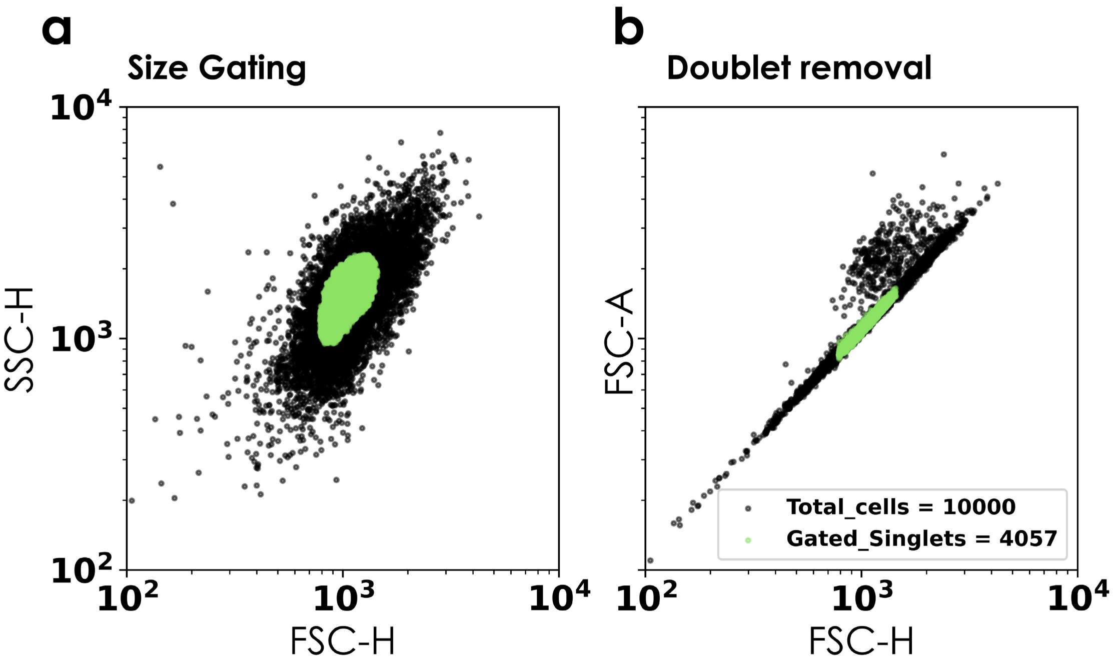


*Figure S1.2. Kernel density based gating of data on size (left) and doublet removal (right). Raw data is shown in black. Gated data is shown in green.*

In cases when multiple fluorescent proteins with overlapping spectra were present inside cells (two recombination cassette strains (Figure 5)), it was difficult to ascertain the differentiation status of the cells. To this effect, we implemented a deconvolution approach previously described in Bertaux et al. 2020[1]. Briefly, 4 single fluorescent protein control strains (mCerulean, mNeonGreen, mVenus, mScarlet-I) with the same promoter and terminator, and integrated in the same locus, were used to determine the spectral signature of each fluorescent protein across the 12 channels of the cytometer (Figure S1.3).

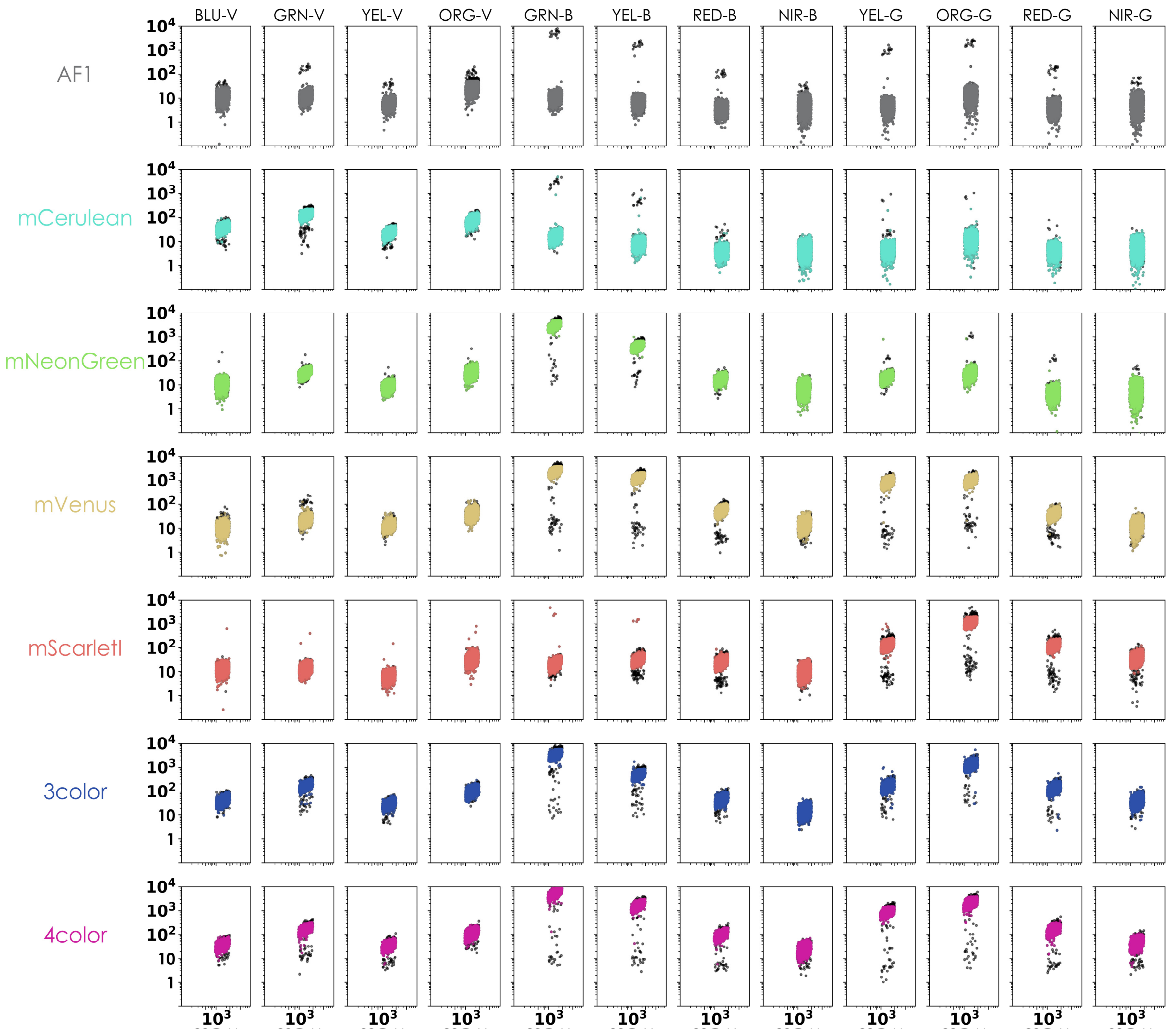


*Figure S1.3. Population distributions of measured intensity by different channels of the flow cytometer for an auto-fluorescence (grey) strain, four single fluorescent protein strains with identical promoters and terminators, mCerulean, mNeonGreen, mVenus and mScarletI, and a four colour strain that has all 4 fluorescent proteins listed above. Data was filtered for outliers using mean absolute deviation metric (4.8 medians) to remove sporadic cross-contamination from other cultures during sampling. We note that filtering was performed only for the experiment used to define spectral signatures.*

These signatures were then used in a linear algebra framework to calculate the individual fluorescence of each fluorophore in a strain harbouring all 4 of the fluorescent proteins. The values for single colour control and the 4 colour strain were in good agreement after deconvolution (Figure S1.4).

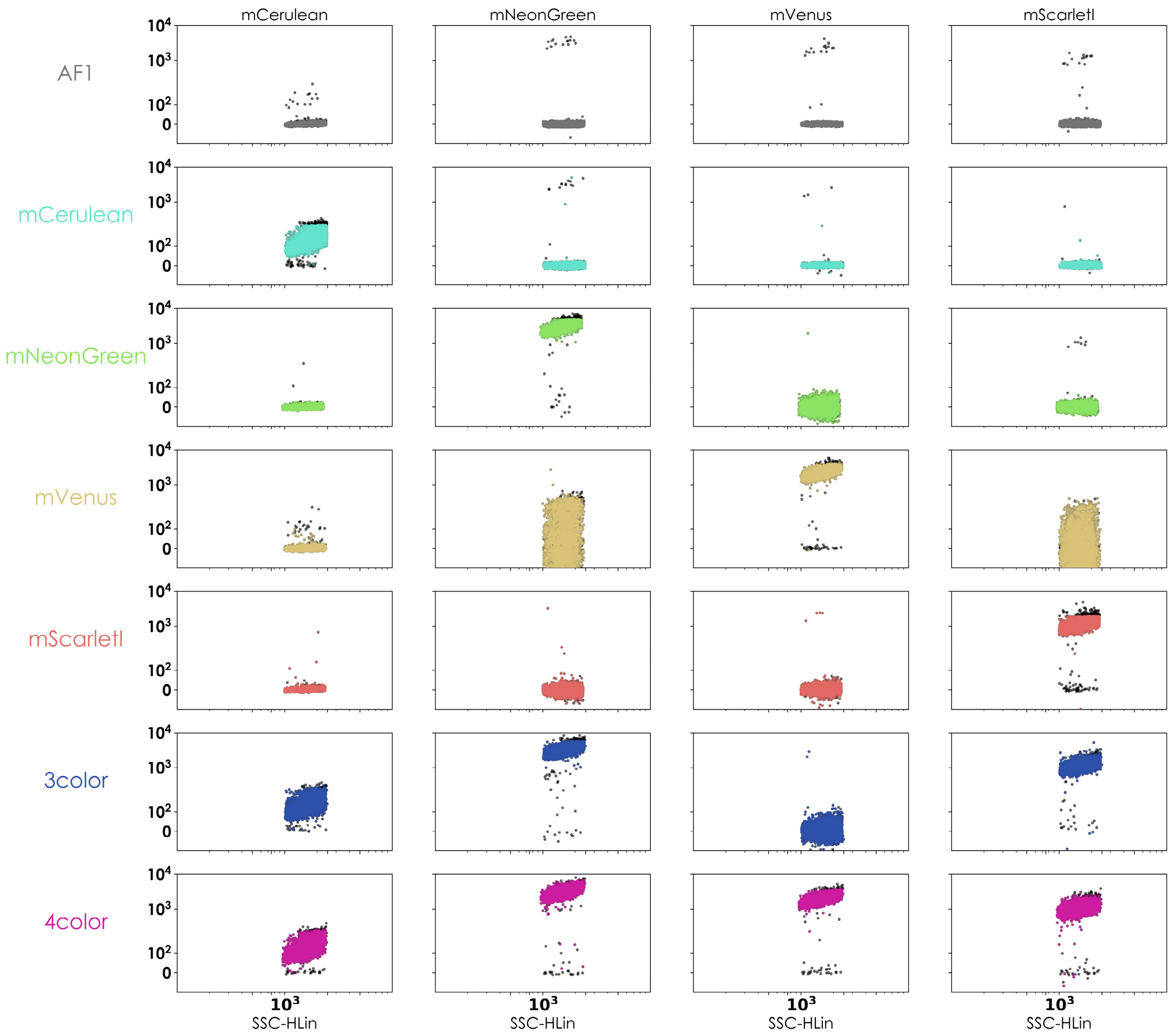


*Figure S1.4. Population distributions of deconvolved fluorescence intensity. Filtered channel data was used to compute spectral signatures of each fluorescent protein. Deconvolution also reduced size related heterogeneity in fluorescence.*

#### II. Microscopy and image analysis pipeline

Microscopy was done on the inverted microscopy platform Leica DMi8 S. Illumination was effected via CoolLED PE-4000 illumination system. Cells were cultured in CellASIC ONIX plate for haploid yeast cells (Y04C) or µIbidi slides.

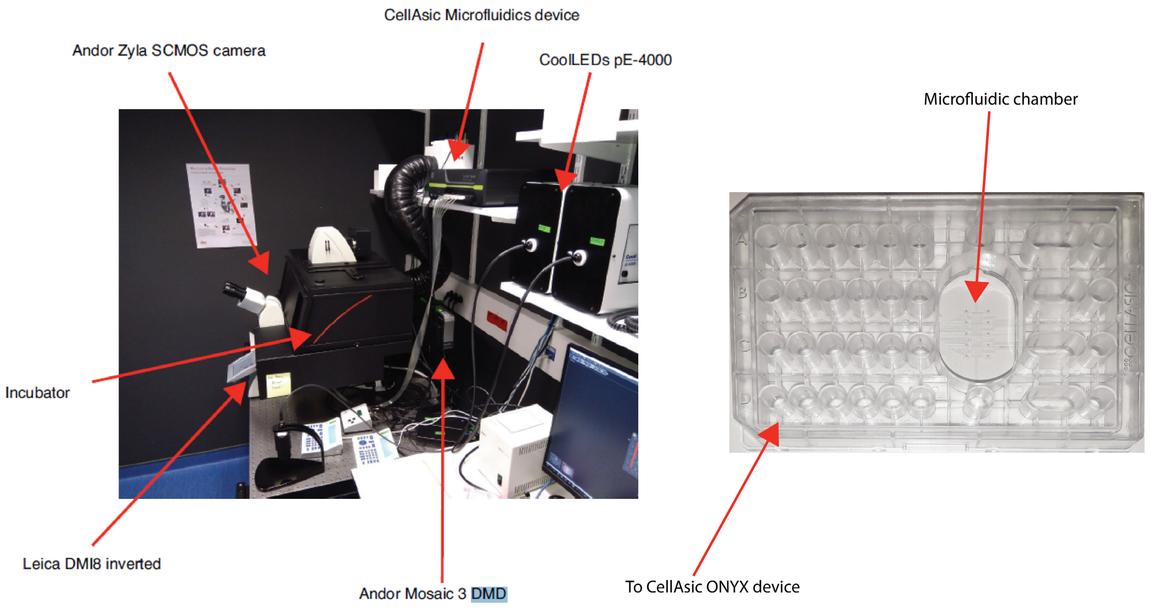


*Figure S2.1. Microscopy platform (left)*[2] *and microfluidic plate (right).*

The spectra of the fluorescent proteins used in this study did not significantly overlap and therefore imaging could be done without the risk of crosstalk or bleed through.

*Table S1. Illumination and imaging conditions.*

| Protein | Filter Cube | Excitation (nm) | Emission (nm) | Dichroic (nm) | Light Source (nm) - Intensity | Exposure (ms) | EL222 Induction |
|---|---|---|---|---|---|---|---|
| mCerulean | CFP | 436/20 | 480/40 | 455 | 435-6% | 600 | Yes |
| mNeonGreen | YFP | 500/20 | 535/30 | 515 | 500-9% | 600 | Yes |
| mScarlet-I | Rhodamine | 546/10 | 585/40 | 560 | 550-10% | 600 | No |

During time-lapse live cell imaging the chamber temperature was maintained at 30°C. We used an in-house software, called MicroMator, for the automated acquisition and cell tracking[2]. Cell segmentation was achieved via SegMator, an in-house U-net based segmentation algorithm[2]. Once cells were segmented, median pixel fluorescence was computed for each cell. Cellular fluorescence was used for ascribing differentiation status to cells. Cells were considered differentiated once a threshold value for fluorescence was exceeded.

We observed that growing in the constrained environment of the microfluidic chamber did not have a significant growth defect (Figure S2.2g).

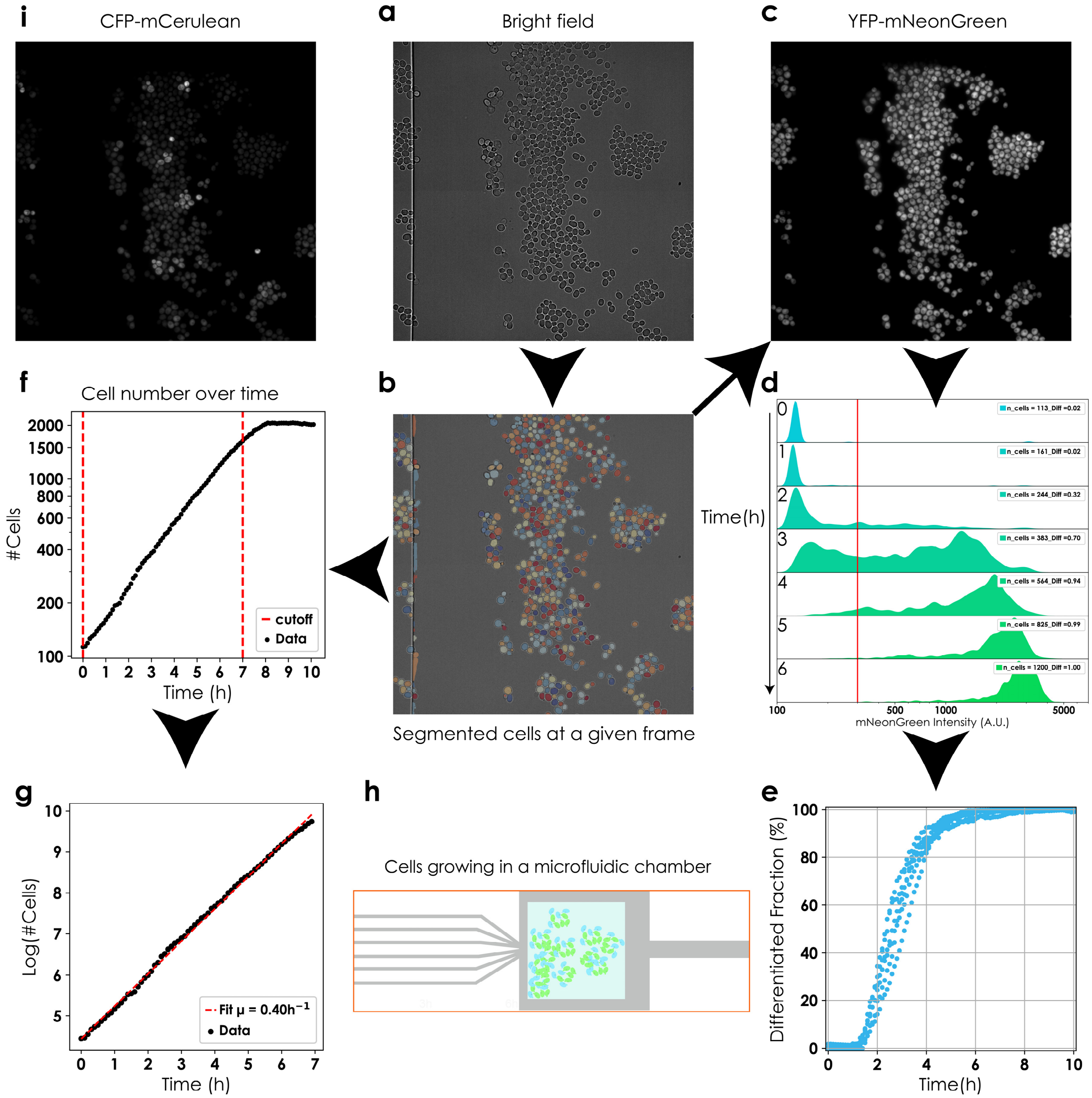


*Figure S2.2. Segmentation and growth in the microfluidic plate. SegMator was used for segmentation and subsequent image analysis[2]. Numbers of cells, over time, in a field of view, were computed by counting the number of segments in each frame. Growth rates were computed by fitting a line to log(#cells) as a function of time. Mean growth rate ± s.d. across 8 fields of view over 2 experiments was found to be 0.394 $h^{-1}$ ± 0.029. Median cellular fluorescence was calculated from segmented images, and fluorescence in the YFP channel was used to distinguish between differentiated and non-differentiated cells. Cells possessing more than 300 units of median fluorescence (A.U.) were considered differentiated. This differentiation fraction was calculated at each frame to illuminate differentiation dynamics (Figure 1**e**).*

#### IIIa. Construction and characterization of the differentiation system

This section is dedicated to the construction and characterization of the differentiation system.

We chose three EL222 promoters published in Benzinger and Khammash, 2018[3] to drive Cre recombinase, namely pEL222 3x binding sites (bs), pEL222 6x bs and pEL222 5x bs Gal1. Of these three, only EL222 5x bs Gal1 led to low leakage with <10% colonies recombined in the dark (data not shown).

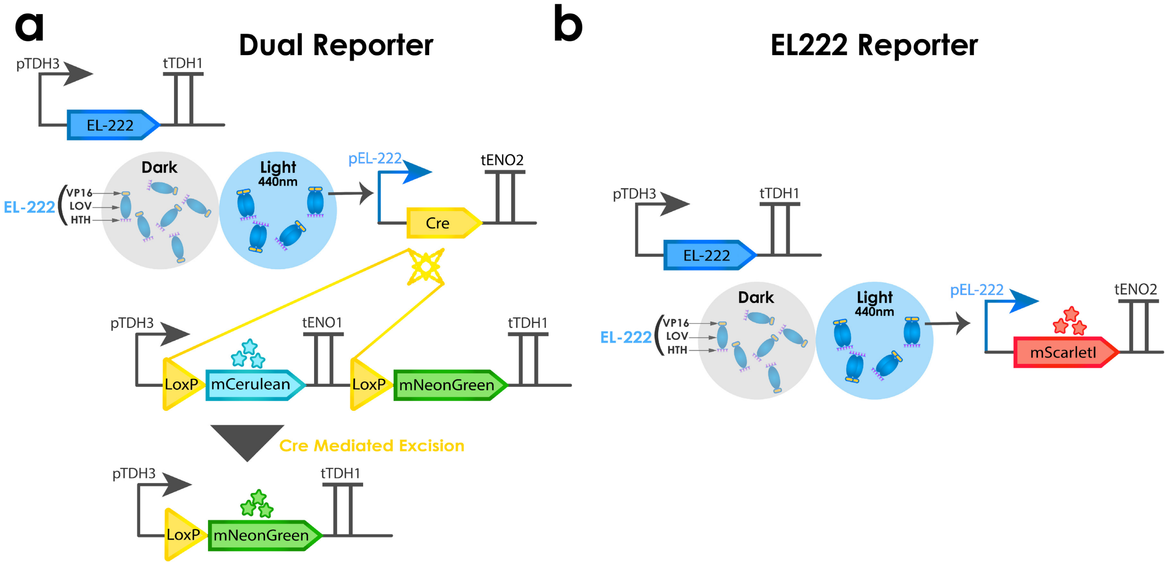


*Figure S3.1. a. Differentiation system (Dual reporter) and b. EL222 activity reporter. ELL222 promoter used in both reporters was pEL222 5X bs Gal1.*

We observed no significant toxicity or growth defect due to expression of EL222, Cre or blue light induction. We also did not notice a growth difference between differentiated and non-differentiated cells.

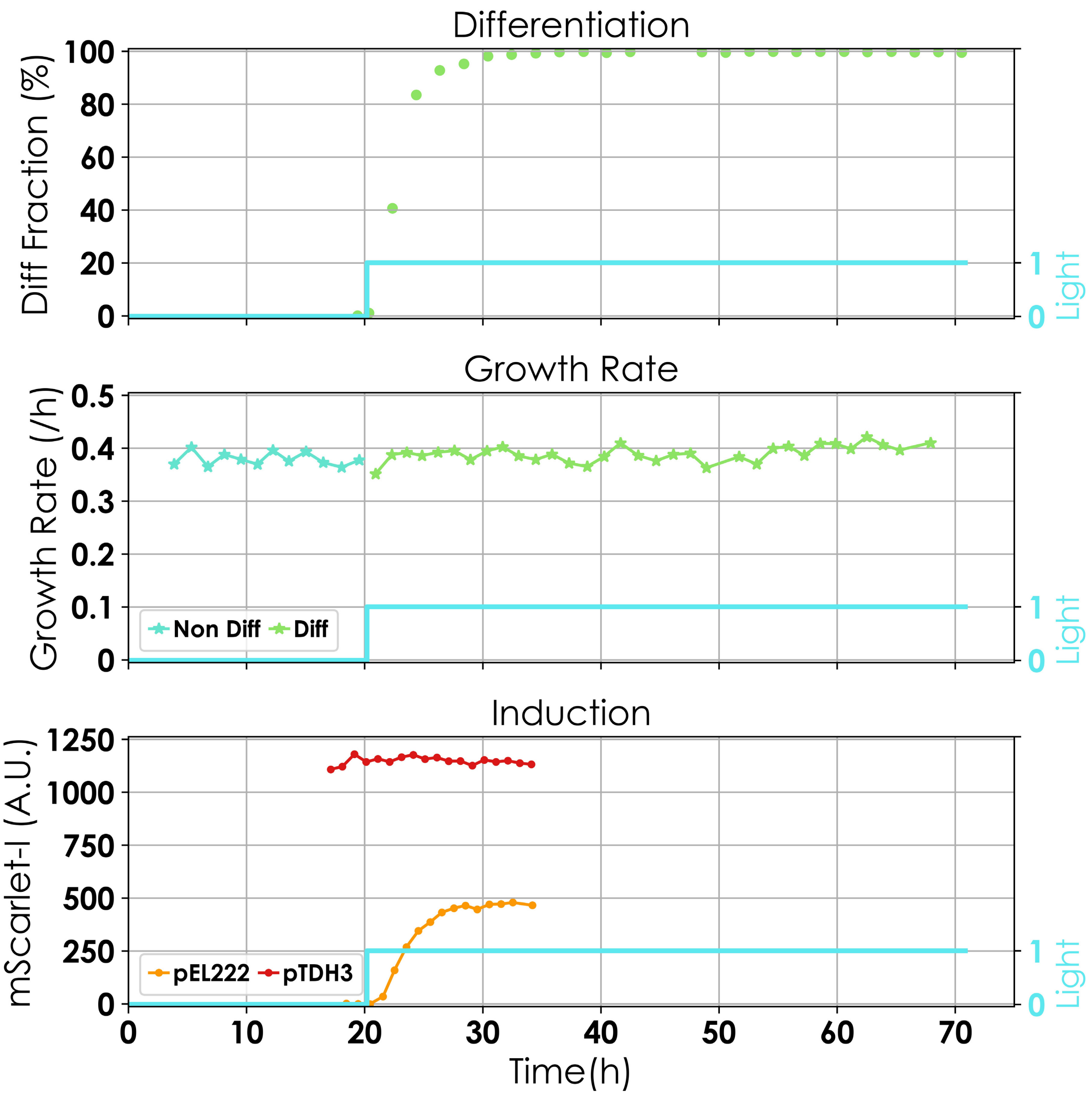


*Figure S3.2 No growth defect is found for differentiated cells, growing under blue light and expressing Cre (top and middle panel). The expression strength of the EL222 promoter is approximately 40% of the one of the TDH3 promoter (bottom).*

Consequently, we decided to study the induction behaviour of this promoter. We put a red fluorescent protein (mScarlet-I) under the control of this promoter (EL222 reporter) and characterized response to various light intensities and duty cycles in cells carrying the EL222 reporter in continuous cultures. By duty cycle is meant a quantity between 0 and 1 that reflects the percentage of light shown in a given period of 2h. For instance, a duty cycle of 0.5 for a period of 2h signifies 1h of light followed by one hour of darkness. Dynamics of population average fluorescence emerging from various different light intensities and duty cycles are displayed in Figure S.3.3a and 3.3b, respectively.

Overall, we obtained results that are consistent with Benzinger and Khammash, 2018[3]:

1. Steady state average fluorescence is linear in the duty cycle and shows a sigmoidal response in increasing intensity of light (Figure S3.3c).
2. Cell-to-cell heterogeneity is larger with intensity modulation for the same mean level of fluorescence compared to duty cycle modulation (Figure S3.3d).

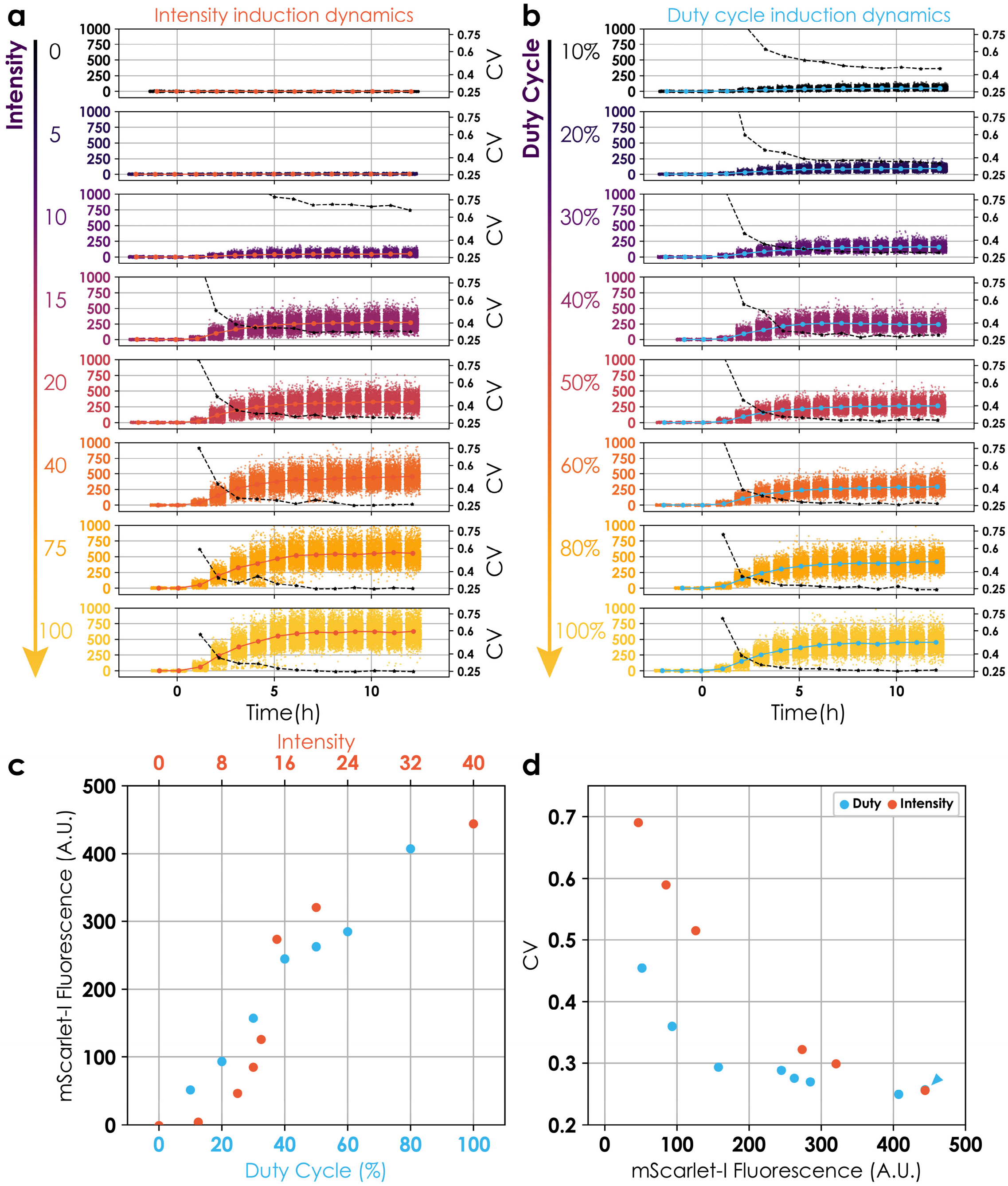


*Figure S3.3. Characterizing EL222 induction behaviour.* ***a*** *and* ***b.*** *Population level induction dynamics for intensity modulation (****a****) and duty cycle modulation (****b****). Small circles represent fluorescence from individual cells. Big circles (red and blue) represent the mean fluorescent values at each time point (connected with red and blue lines, respectively). Black lines marked with stars denotes the CV (measured on the right y-axis). CV values prior to induction were not reliable due to low average population fluorescence.* ***c.*** *Red circles represent mean population fluorescence for duty cycle modulation and blue circles stand for mean population fluorescence for intensity modulation. Each condition has a single replicate.* ***d.*** *CV plotted against mean for intensity modulation (red) and duty cycle modulation (blue).*

Next, we tested the differentiation system for different intensities and duty cycles in exponentially growing cells in batch in duplicates. These experiments were done in batch because it allowed us to perform multiple induction experiments on the same day. Briefly, precultures could be diluted in a large volume and partially split to give 16 cultures in falcon tubes that could be induced with different profiles and removed from the batch inducer and placed in the dark while the same preculture could be used to conduct the next set of experiments. The resulting differentiation fractions (in duplicates) were reminiscent of EL222 induction profiles.

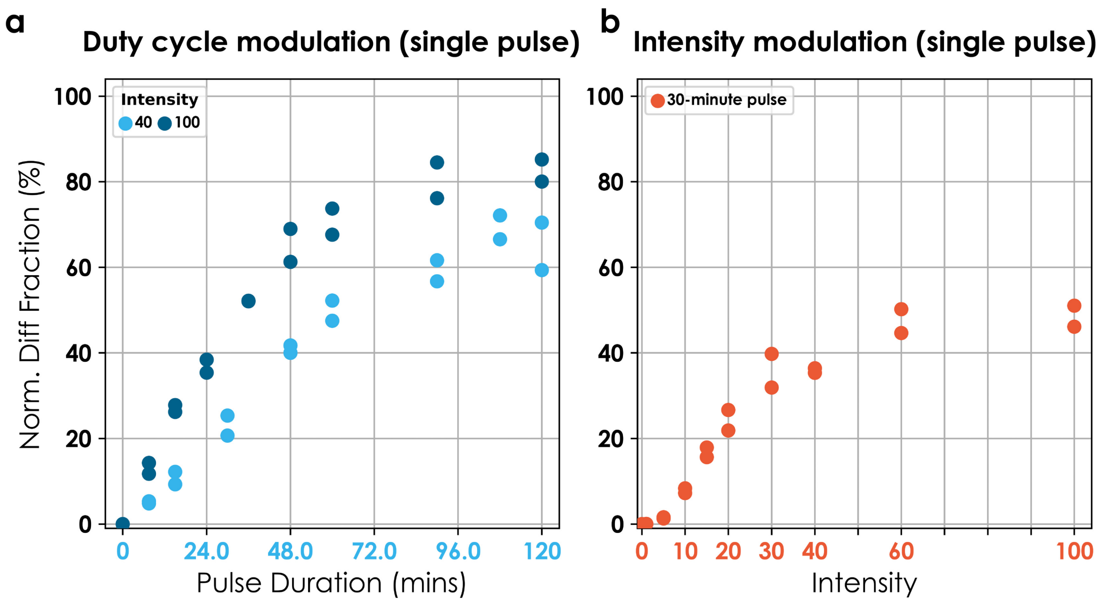


*Figure S3.4. Single pulse duration vs intensity modulation of differentiation behaviour. Solid circles are differentiation fractions observed. All conditions have two replicates. Differentiation w.r.t. pulse duration modulation over a period of 2h was assessed for two different intensities, 40 and 100. We noticed that the differentiation fraction increases linearly with pulse duration up to a certain point whence it plateaus. The slope of increase in differentiated fraction with respect to pulse duration was steeper for intensity 100. The linear response range was also smaller for intensity 100. Amplitude modulation led to a more switch like response of the differentiated fraction. Intensity 10 resulted in no significant recombination whereas intensity 30 resulted in 80% differentiation of the maximum intensity (100).*

For all experiments in the main text, an intensity of 40 was used.

We compared the efficiency of our system with existing systems of optogenetic recombination in yeast. We found that our system was the most efficient light based recombination system to date. The table compares the efficiency in yeast. Since, the systems reported in Taslimi et al. 2016[4] and Kawano et al. 2016[5] consisted of results for mammalian cells (HEK293 and COS-7, respectively), data for these two is taken from Duplus-Bottin et al. 2021[6]. Leakage in dark is defined as percentage of cells identified as recombined after culture in the dark.

**Table S2. Characteristics of optoinducible/photoactivable recombinases in yeast[4–7].**

| **Publication** | **System** | **Growth conditions** | **Efficiency (mean ± s.d.) - Induction time** | **Leakage in dark (mean ± s.d.)** |
|---|---|---|---|---|
| Ref 6 (Figure 3), adapted from Ref 4† | split Cre CIB1_CRY2 | Stationary liquid culture | 1.6% ± 0.8 at 90 mins | 1.3% ± 0.5 over 24h of dark culture |
| Ref 6(Figure 3), adapted from Ref 5†† | split Cre pMag-nMag | Stationary liquid culture | 21.2% ± 5.8 at 90 mins | 7.1% ± 1.1 over 24h of dark culture |
| Ref 7 (Figure 2) original study | split Cre PhyB-PIF3* | Stationary liquid culture | 46.7% ± 5.3 at 24 h** | 6.7% ± 2.6 over 24h of dark culture |
| Ref 6 (Figure 3 & 4) original study | destabilized Cre fused to asLOV2 (LiCre) | Stationary liquid culture<br>Exponential liquid culture | 41.2% ± 2.8 at 40 mins<br>66.7% ± 3.7 at 90 mins<br>66.8% ± 3.3 at 180 mins<br>7.6% ± 2.1 at 40 mins | 0.7% ± 0.2 over 24h of dark culture |
| This study | EL222 inducible WT Cre | Exponential liquid culture | **43.1% ± 2.7** at 40 mins<br>**76.8% ± 1.7** at 90 mins<br>**94.4% ± 0.9** at 180 mins<br>**99.7% ± 0.1** at 240 mins | **0.06% ± 0.05 over 72h of dark culture** |

† Original study performed in HEK293 cells.
†† Original study performed in COS-7 cells.
*Addition of PCB was necessary to achieve recombination with PhyB-PIF3 system. Results shown here correspond to a concentration of 25µM. Efficiency could be improved to 89.3% ± 4 by increasing the concentration of PCB to 100µM. This also resulted in an increased the background activity to 9.7% ± 2.4.
** Induction was carried out by delivering a single 5-min pulse of red light and followed by 10s pulses every 5 minutes for 24h.

#### IIIb. Model Development

We developed a model to assess whether our system is predictable i.e. whether we could predict the dynamics of the differentiated fraction emerging from given light inputs. Since the model was primarily developed for the purpose of deploying it in model predictive control of the population composition, we decided to use a simple ODE model that tracks population dynamics.

$$\dot{g}(t) = \mu g(t) - \boldsymbol{U}(t) r_{diff} g(t) - \lambda g(t)$$

$$\dot{p}(t) = \mu p(t) + \boldsymbol{U}(t) r_{diff} g(t) - \lambda p(t)$$

$$n = g(t) + p(t)$$

$$\lambda = \mu$$

$g$ and $p$ stand for specific cell density (in O.D. units) of non-differentiated and differentiated cells, respectively. μ is growth rate per hour of the culture. $\boldsymbol{U}(t)$ is light signal as a function of time and can take values 0 or 1. $r_{diff}$ is the differentiation rate under continuous light. $n$ is the total cell density (in O.D. units) of the culture and is kept constant. $\lambda$ is the dilution rate. At constant total cell density, dilution rate, $\lambda$ equals growth rate, μ. No significant difference was observed in the growth rate of differentiated cells and non-differentiated cells (Figure S3.2).

The ODE model was conceived with only one free parameter, namely the differentiation rate $r_{diff}$. Initial conditions and the growth rate were fixed from the data (Figure 2d). Experiments were performed in the bioreactor platform, which showed modest reactor-to-reactor variability.

Data fitting was implemented in Python using SciPy library. Bounds were imposed on the parameter search ($10^{-10}$ -10) to preserve physiological relevance. Parameter searches were conducted locally but starting from different initial parameter values spanning four orders of magnitude using an approach similar to Branch, Coleman and Li[8]. In all cases, parameter searches converged to the same parameter value. Our results suggest that this simplistic model can be used to predict steady state differentiation levels given an input light signal. The model could well predict the differentiation dynamics if we assume a delay of 60 minutes to account for observation delays that were not explicitly included in the model. (Figure 2g).

#### IVa. Construction and characterization of GAuDi strain

This section is dedicated to the construction and characterization of the differentiation system coupled to a growth arrest module such that cells growth arrest upon differentiation.

In order to arrest the cells upon differentiation, we hijacked the mating factor pathway by overexpressing a variant of the downstream effector FAR1. Concretely, prior to recombination, cells express mNeonGreen. After recombination, cells start expressing ATAF1, an orthogonal transcription factor, that drives expression of FAR1M[9] (growth arrest) and mScarlet-I (classification) with 4x binding site promoter[10]. A positive feedback loop was introduced to increase the levels of ATAF1 and consequently, FAR1M. Concretely, we added a transcriptional unit with pATAF1 driving ATAF1 TF. The strain will be referred to as Gaudi in the following (Figure S4.1).

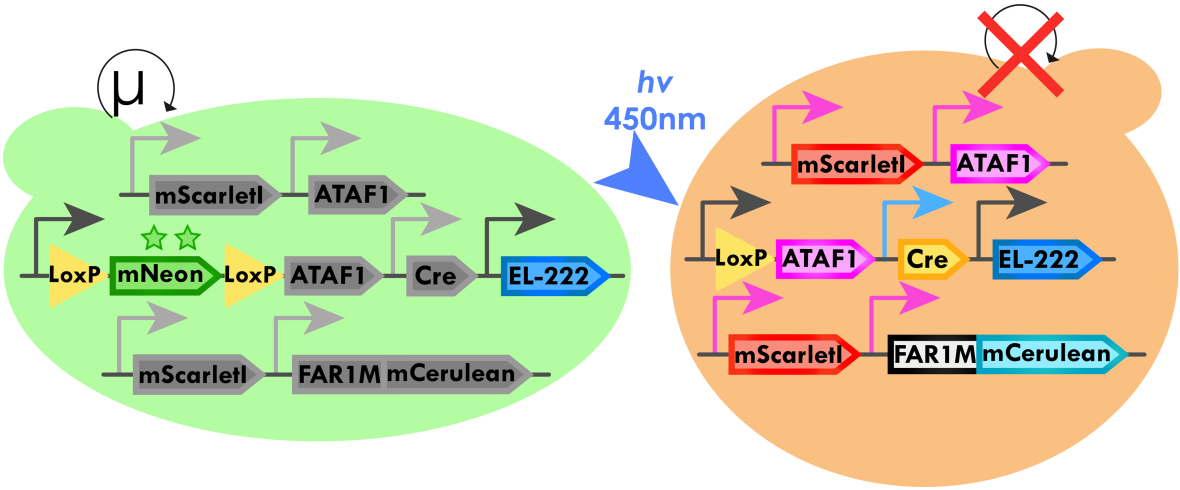


*Figure S4.1. Circuit for GauDi system. We added a positive feedback loop with ATAF1 TF driving its own expression to ameliorate the expression levels of FAR1M.*

We conducted experiments in turbidostat and under the microscope in a microfluidic chamber and found that this topology enabled growth arrest upon differentiation. Under the microscope, cells did not undergo any cell divisions after light application.

We grew cells in the turbidostat with continuous light and estimated the growth rate by measuring the OD (Figure S4.2). We noticed that growth rate came to a halt roughly 4 hours and the arrest lasted for 15-20h post light induction and then the cultures escaped. Along with the OD measurements after light induction, we also analysed the population composition and discovered a variable, but consistently present, population of cells that contained neither green nor red. Presumably, these cells are dead cells.

We suspected that the escapers were mutants and extracted the genomes of 5 escapers and sent them for targeted genome sequencing. Results suggested that loss of the integrative cassette was the most common mutation (3/5) that led to escape. We hypothesized that if this were the case, mutants that lose the entire integrative cassette, including the auxotrophic marker, would be selectively disadvantaged. Therefore, we cultured cells under continuous stimulation but this time in selective media. We observed that the cells could be arrested for 50-100% longer, suggesting that other mutations could also lead to escape. Curiously, the growth rate of differentiated cells was higher in selective media (Figure S4.2).

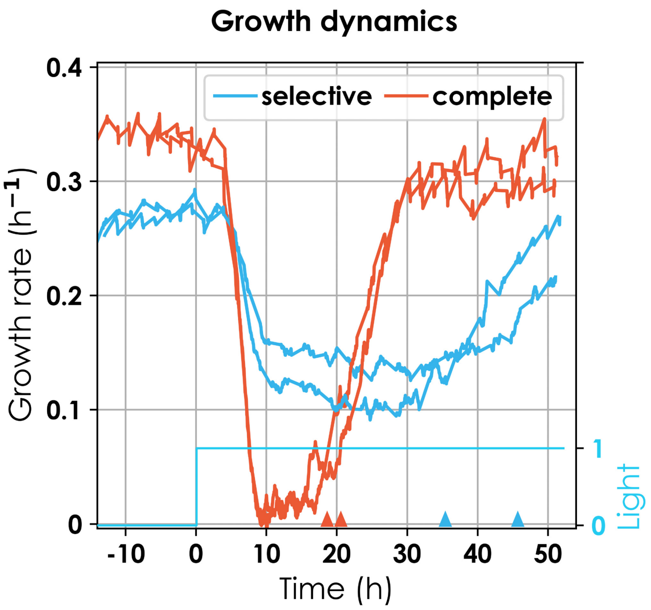


*Figure S4.2. Growth rate of GAuDi02 under continuous light in non-selective (red) and selective (blue) media in duplicates. The cyan line signifies light induction. Manually placed triangles indicate the time when the culture as a whole escaped the growth arrest.*

Next, we characterized the differentiation dynamics of the strain in response to light. We applied different light pulses at different intervals to cultures. We observed a fast decrease in the differentiated fraction once light was removed suggesting that the growth arrest was strong and that this system could be used for dynamic control of microbial consortium in a self-contained configuration.

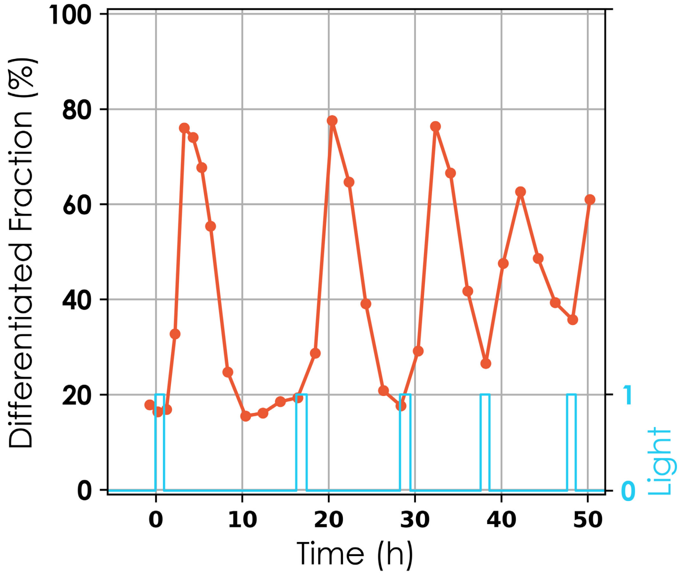


*Figure S4.3. Differentiation dynamics (red-dotted line) for periodic light pulses (cyan line). We observed that light pulses led to increases in differentiated fractions that subsided rapidly after removal of light (a, b).*

#### IVb. GAuDi model development

Based on characterization data, we introduced two new state variables in the ODE model, escapers and dead cells. The rate of escape and the death rate could not be directly fixed from the data and are treated as unknown model parameters. Furthermore, due to the leakiness of the ATAF1 feedback loop, we observed a persistent fraction of cells that appeared to be initially differentiated given their levels of mScarletI fluorescence. Since empirically it was not possible to separate the differentiated cells from those with an overactive feedback loop, we introduced a basal differentiation rate present in the absence of light (and its presence). The growth rates of differentiated, non-differentiated and escapers were fixed from the data.

$$\dot{g}(t) = \mu_g g(t) - \left(\mathring{r}_{diff} + \boldsymbol{U}(t) r_{diff}\right) g(t) - \lambda g(t)$$

$$\dot{p}(t) = \mu_{diff} p(t) + \left(\mathring{r}_{diff} + \boldsymbol{U}(t) r_{diff}\right) g(t) - r_{esc} p(t) - r_{dead} p(t) - \lambda p(t)$$

$$\dot{e}(t) = \mu_{esc} e(t) + r_{esc} p(t) - \lambda e(t)$$

$$\dot{d}(t) = r_{dead} p(t) - \lambda d(t)$$

$$n = g(t) + p(t) + e(t) + d(t)$$

$$\lambda = \frac{g(t)}{n} \mu_g + \frac{p(t)}{n} \mu_{diff} + \frac{e(t)}{n} \mu_{esc}$$

$g$, $p$, $e$, and $d$ are specific cell density (in O.D. units) of non-differentiated, differentiated, escaper, and dead cells, respectively. $\mu_g$, $\mu_{diff}$, and $\mu_{esc}$ are growth rates per hour of non-differentiated, differentiated, and escaper cells, respectively. $r^{\circ}_{diff}$ is basal differentiation rate and $r_{diff}$ is the differentiation rate per hour in presence of light. $\boldsymbol{U}(t)$ is the light signal as a function of time and can take values 0 or 1. $r_{esc}$ is the rate of escape per hour from growth arrest in differentiated cells per hour. $r_{dead}$ is the rate of death per hour in growth arrested differentiated cells per hour. $n$ is the total cell density (in O.D. units) of the culture. $\lambda$ is the culture dilution rate and, at constant cell density, equals the weighted sum of growth rates of individual species.

The model was fitted to dynamical data (differentiated & dead fraction) with non-trivial light signals (Figure 4b). Concretely, the differentiation fraction and the dead fraction were computed from time-series flow cytometry data to fit the model. However, there were significant delays between differentiation events and the time by which corresponding cells could be classified as differentiated based on their fluorescence. Similarly, differentiated cells died only after being arrested for some time and the growth arrest required two cell generations to fully manifest. To account for these delays, model predictions were shifted in time (2h for differentiated and 6h for dead cells, and 5h for the growth rate). Bounds were imposed on the parameter values ($10^{-10}$ -10) to preserve physiological relevance. The parameter search was conducted locally but starting from different initial parameter values spanning four orders of magnitude using an approach similar to Branch Coleman and Li[8]. In all cases, the search converged to the same parameter values.

| ***Table S3*** | *Parameter estimates* (/h) *for ODE model for GAuDi02.* | | | | | | |
|---|---|---|---|---|---|---|---|
| Parameter | $\mu_g$ | $\mu_{diff}$ | $\mu_{esc}$ | $\mathring{r}_{diff}$ | $r_{diff}$ | $r_{esc}$ | $r_{dead}$ |
| Estimated Value | 3.9e-01 | 4.0e-02 | 3.9e-01 | 6.98e-02 | 1.16e+00 | 9.43e-08 | 7.12e-02 |

Parameters in green in table S3 were fixed directly from the data while those in yellow were e stimated.

These parameters led to good agreement between model outputs and observed data (Figure 4b). We further validated the model by predicting the outcome of four other experiments with dynamic non-trivial light inputs. Model predictions were in good agreement with observed data (Figure S4.4). However, differentiation dynamics and growth arrest could not be predicted well under continuous light.

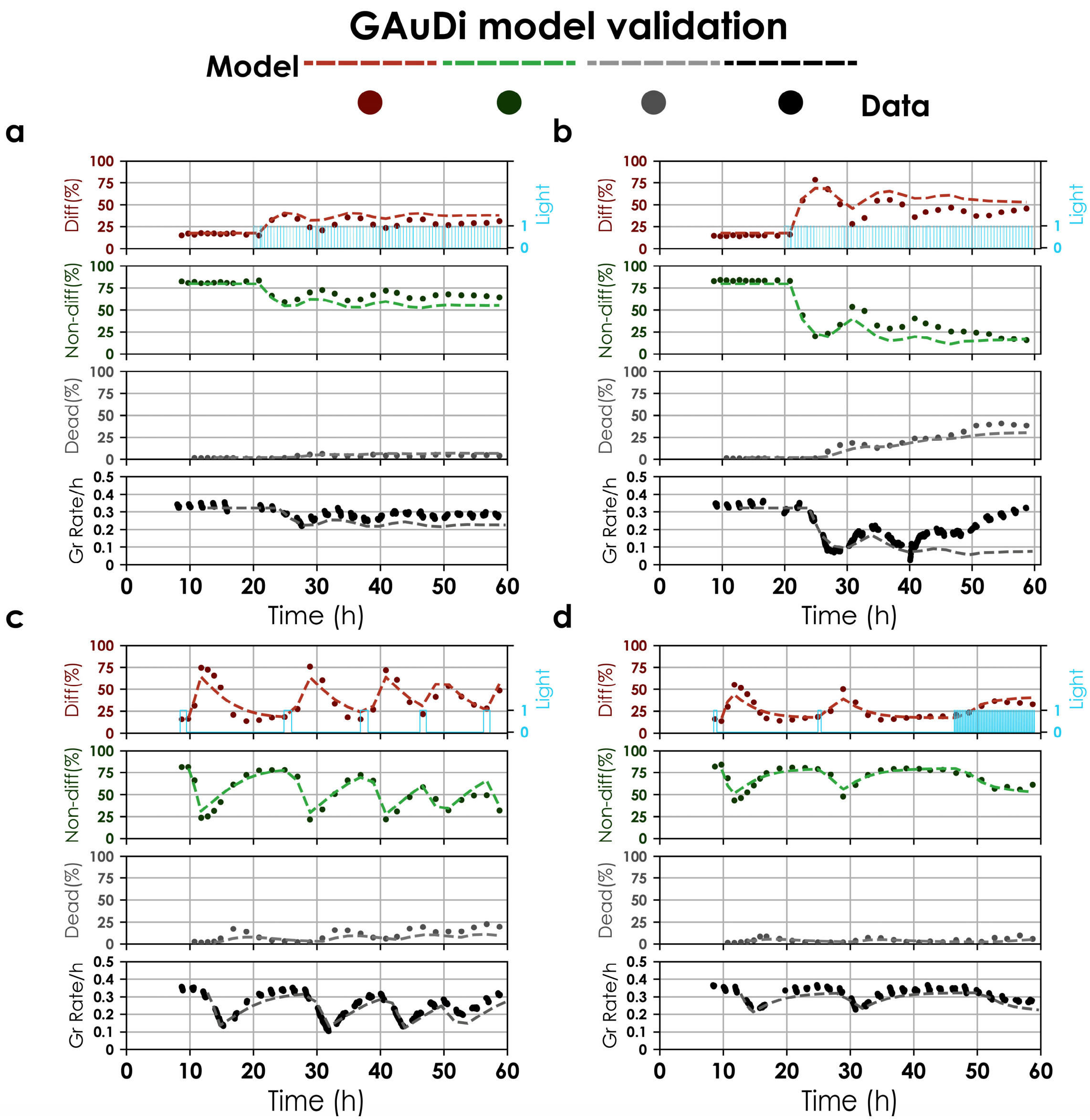


Figure S4.4. *GAuDi model predictions and data for 4 light profiles. The fitted model was validated by using it to predict dynamic responses to light signals that were not used for the fit (blue lines). Circles represent differentiation fractions from a single experiment and dotted lines represent model predictions. The red line represents differentiated fraction, green is the non-differentiated fraction, grey is the dead fraction and black is the culture growth rate. Model predictions were shifted in time to account for observation delays. Initial conditions were fixed from the data.*

#### V. MPC Experiments

##### General strategy

1. Flow cytometry data is processed online to yield state estimates.
2. Some variables, like the escaper fraction, are non-measurable and therefore estimated from the model.
3. Conversion of this state estimate into a real-time state estimate.
   a. Because of delay not captured in the model, state estimates are taken to correspond to;
   $T_0$ = sampling time – (sampling delay + observation delay)
      i. Sampling delay was the time between sample was taken and the data acquisition finished.
      ii. Observation delay was the delay between change of genetic state of a cell and by the time enough fluorescence was produced to classify it as differentiated cell.
   b. Forward simulation of the model to go from $T_0$ to $T_{now}$ (given the actual light sequence applied during that time window)
4. Optimization of the next sequence of 10 duty cycles (horizon = 5h) so as to minimize the predicted error between target and model prediction (gradient descent, scipy.optimize)

**Dynamic control and ODE model adjustments**

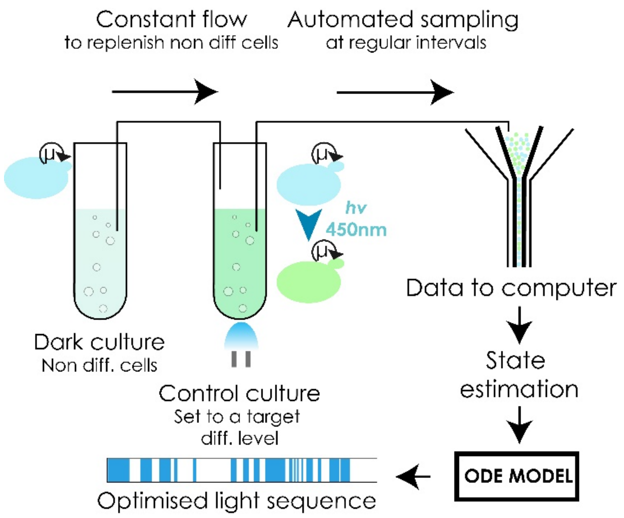


*Figure S5.1. Experimental setup.*

Both reactors were maintained at constant ODs but different from each other. We wondered how to set the ODs of the two reactors so as to ensure that up to 90% of cells could be differentiated in the control reactor. Since there was a constant flux of non-differentiated cells, a flux term had to be added in the ODE model. We also observed that growth rates of cultures at different ODs were not equal. We assume, however, that cells that enter the control reactor from the reservoir reactor instantaneusly change their growth rate.

$$\frac{dg}{dt} = \mu_{target} g - \boldsymbol{U}(t)\, r_{diff} g + Q - \lambda g$$

$$\frac{dp}{dt} = \mu_{target} p + \boldsymbol{U}(t)\, r_{diff} g - \lambda p$$

$$Q = \mu_{reservoir} n_{reservoir}$$

$$n_{target} = g + p$$

$g$ and $p$ stand for the specific cell density (in O.D. units) of non-differentiated and differentiated cells, respectively. $\mu_{control}$ and $\mu_{reservoir}$ are growth rates in the control and reservoir reactors, respectively. $\boldsymbol{U(t)}$ is the light signal as a function of time and can take values 0 or 1. $n_{target}$ and $n_{reservoir}$ are O.D. ( total cell densities) at which control and reservoir reactors are maintained, respectively. $r_{diff}$ is the differentiation rate under continuous light. $Q$ is the flux of non-differentiated cells from reservoir to control reactor. $\lambda$ is the dilution rate of the control reactor.

Since we maintain the control reactor at constant OD, the number of cells in the control reactor stays constant at steady state.

$$\frac{dn_{target}}{dt} = 0$$

$$\frac{d(g+p)}{dt} = 0$$

$$\frac{dg}{dt} + \frac{dp}{dt} = \mu_{target}(g+p) + Q - \lambda(g+p)$$

$$0 = \mu_{target}(g+p) + Q - \lambda(g+p)$$

$$\lambda = \mu_{target} + \mu_{reservoir} \frac{n_{reservoir}}{g+p}$$

$$\lambda = \mu_{target} + \mu_{reservoir} \frac{n_{reservoir}}{n_{target}}$$

$$\lambda = \mu_{target} + \alpha\mu_{reservoir}$$

α is the ratio of reservoir OD to control reactor OD.

Let $r$ be equal to the fraction of differentiated cells in the control reactor.

$$r = \frac{p}{g+p}$$

$$\frac{dr}{dt} = \frac{\frac{dp}{dt}}{g+p}$$

$$\frac{dr}{dt} = \frac{\mu_{target}p + \boldsymbol{U}(t)r_{diff}g - (\mu_{target} + \alpha\mu_{reservoir})p}{g+p}$$

$$\frac{dr}{dt} = \boldsymbol{U}(t)r_{diff}(1-r) - \alpha\mu_{reservoir}r$$

$$\frac{dr}{dt} = -(\boldsymbol{U}(t)r_{diff} + \alpha\mu_{reservoir})r + \boldsymbol{U}(t)r_{diff}$$

This is a first order non-homogeneous linear differential equation in standard form,

$$\frac{dy}{dt} = -p(t)y + f(t)$$

For maximum differentiation fraction achievable ($\boldsymbol{U}(t) = 1$), at steady state,

$$\frac{dr}{dt} = 0$$

$$r_{diff} = (r_{diff} + \alpha\mu_{reservoir})r_{max}$$

$$\alpha = \frac{r_{diff}}{\mu_{reservoir}}\left(\frac{1}{r_{max}} - 1\right)$$

Plugging in values of $r_{diff}$ = 0.86 $h^{-1}$, $r_{max}$ = 0.9 and $\mu_{reservoir}$ = 0.41 $h^{-1}$ we get, $\alpha$ = 0.23.

Therefore, to reach a target differentiation level of 90 % in the control reactor, the OD of the reservoir had to be ~5 times lower. Based on this calculation, the OD of the reservoir reactor was set to 0.1 and that of control reactor to 0.6. Growth rates were determined from the data. This model was then plugged into the MPC framework described previously in Bertaux et al. 2020[1]. Briefly, the MPC algorithm utilizes the fitted model to search for the optimal sequence of light signals that would lead to desired target level of differentiated cells. The differential equation obtained in (2) could be solved analytically for constant light signal and was solved piecewise in the model. Receding time horizon for model predictions for the experiments shown here was 5 hours (10 cycles of period 30 minutes each). Automated flow cytometry measurements were performed every hour. The light sequence was automatically re-optimized and updated at every timepoint based on the newly estimated state of the system. There existed a significant delay (1.5-2h) between the measurement and the state of the system at the time of measurements. To bridge this gap, state estimation was used. We assumed that the measured data corresponds to a snapshot of the population composition 1.5h in the past and to estimate the current state we solved equation (2) with the light sequence that was applied over the past 1.5h. This was critical to avoid oscillatory behaviour of the system. Furthermore, we noticed that the differentiation rate parameter estimated from experiments led to a steady state error. Therefore, the value of this parameter was reduced to 0.4 /h in the MPC model to ensure robust control.

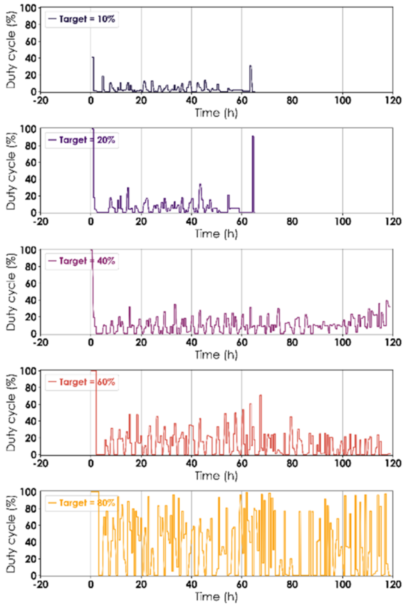


*Figure S5.2 Light profiles for experiments in Figure 3b. Duty cycles were estimated for a period of 30 minutes. Induction started at t=0.*

**Single vessel control**

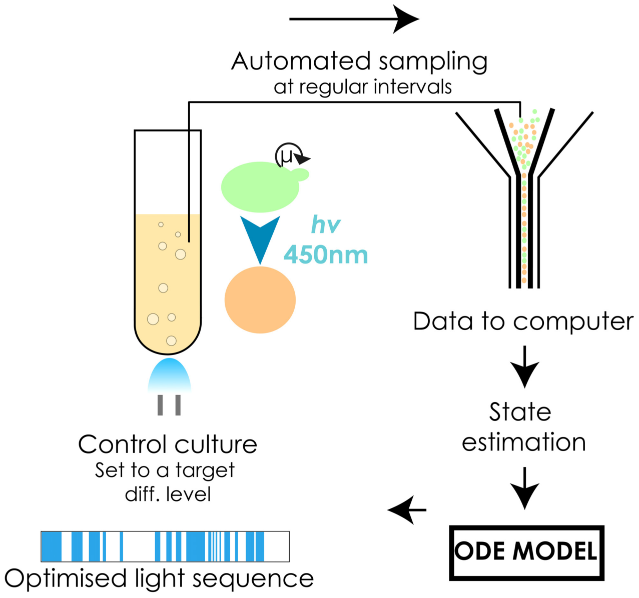


*Figure S5.3. Experimental setup.*

The model for the GAuDi strain was used for the MPC. No adaptation to the model was necessary. We could only sample the culture every two hours owing to volume limitations and consequently the delay was increased to 2.5 hours.

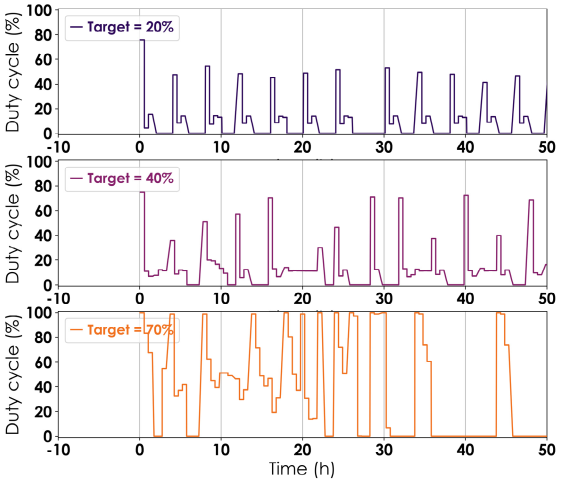

*Figure S5.4 Light profiles for experiments in Figure 3e. Duty cycles were estimated for a period of 30 minutes. Induction started at t=0.*

### VI. Spatial control

Cells harbouring the differentiation system were cultured in the dark to late exponential phase in 2% glucose synthetic complete LoFlo media at 30°C in 50mL falcon tubes. Cells were then loaded in a 6 channel µIbidi slide (Catalogue # ;3939) such that they formed a monolayer. The µIbidi slide was induced under the microscope (10x magnification) within a user-defined pattern with the help of a digital mirror device (Andor Mosaic3). The pattern was a binary bitmap with black representing no light, and white representing induction.

A light of 460 nm was used for induction with the CFP band pass filter (see Table S2) . The induction was effected for 1s at intensity 10% every 3 minutes for 1h. Following the induction, cells were allowed to recover for 1h prior to imaging. This also gave time for enough mNeonGreen molecules to mature and be detected. It was difficult to increase the duration of induction period, even though light was not lethal for cells, because of cell growth, and subsequent emergence of bubbles displacing the monolayer. We tried to ameliorate the growing conditions by culturing cells in a microfluidic plate but the media flow disallowed any meaningful quantification of efficiency of pattern formation at low cell density. This was because buds from inside the pattern would be transported with the flow, to regions outside the pattern. When we tried to do the experiment after cells had formed a monolayer, flow was obstructed leading to growth rate decrease and cell death due to lack of media. We note that the cells could not be imaged during the induction period as both mCerulean and mNeonGreen required illumination that would activate the EL222 and lead to undesirable differentiation.

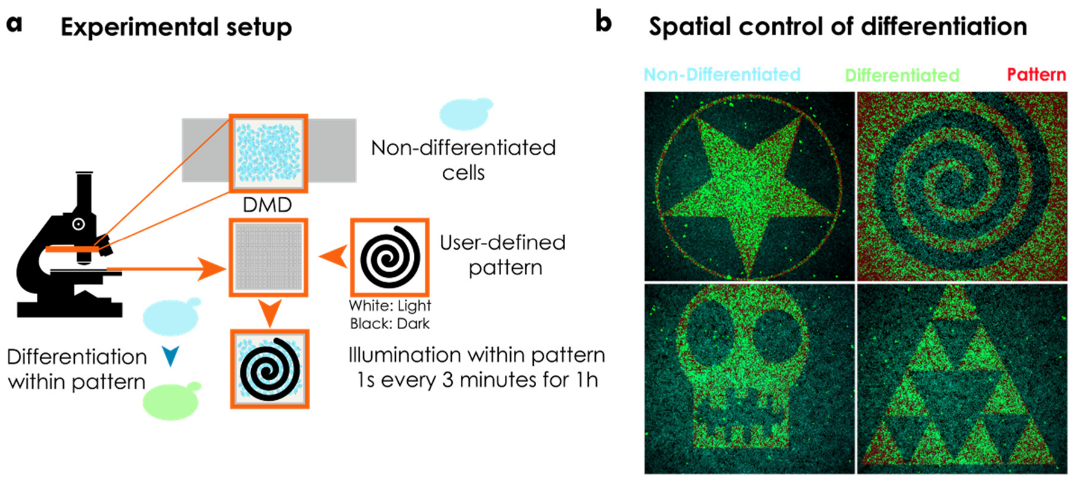


*Figure S6.1 Precise spatial control of heterogeneous microbial consortia originating from a single strain. a. Hardware setup for realizing pattern formation under the microscope. Cells carrying the differentiation system were cultured in the dark until late exponential phase (OD ~0.8) and loaded in a µIbidi slide. Cells were allowed to settle down and form a monolayer. Light was shone in a user-defined pattern over the monolayer using a digital mirror device (DMD). Light was shone as 1s pulses every 3 minutes for 1h (20 pulses). Following light delivery, cells were kept in darkness for 1 hour before imaging them. b. Images of imprinted patterns in the population. mCerulean fluorescence was ascribed a cyan colour, whereas mNeonGreen fluorescence was ascribed a green colour (false colour in microscopy images). The pattern, in red, is overlaid on top of the merge for cyan and green channels.*

#### VII. Expanding the differentiation system to give rise to multi-species consortia with differentiation programs

In order to explore the possibility of engineering differentiation programs, we cloned two recombination cassettes of unequal to-be-excised region but otherwise identical, in different strains. These strains were cultured continuously in the turbidostat and induced with full light. We found that changing the length of the to-be-excised region plays a crucial role in determining the rate of differentiation (*Figure S7.1*). We observed that differentiation rate is inversely proportional to the length of the to-be-excised region.

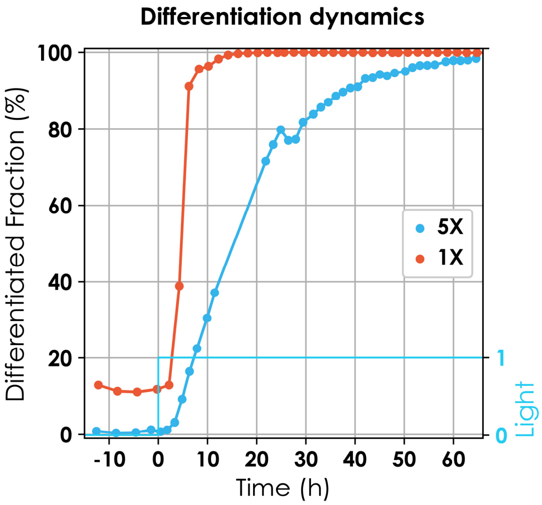


*Figure S7.1. Differentiation fraction dynamics for recombination cassettes of different to-be-excised regions (**1X**:red and **5X**:blue) in continuous light. Red and blue dotted lines show differentiation fraction over time. Legend indicates the length of the to-be-excised region compared to the one used for Figure 1. For the experiment shown here, a significant part of the population for **1X** was recombined prior to light induction.*

Exploiting this information, we implemented differentiation programs by introducing two recombination cassettes of to-be-excised-regions of different lengths. These were named according to their differentiation behaviour with respect to the two recombination cassettes. If the recombination cassettes are of equal length (*Figure S7.2a*), the recombination is asynchronous meaning either recombination cassette could recombine independently of each other (*Figure S7.3a*). We can achieve sequential differentiation if the to-be-excised regions are unequal (*Figure S7.2b*) meaning that the recombination occurs first at the shorter site followed by recombination at the longer site. We did not observe recombination of the longer site exclusively (*Figure S7.3b*).

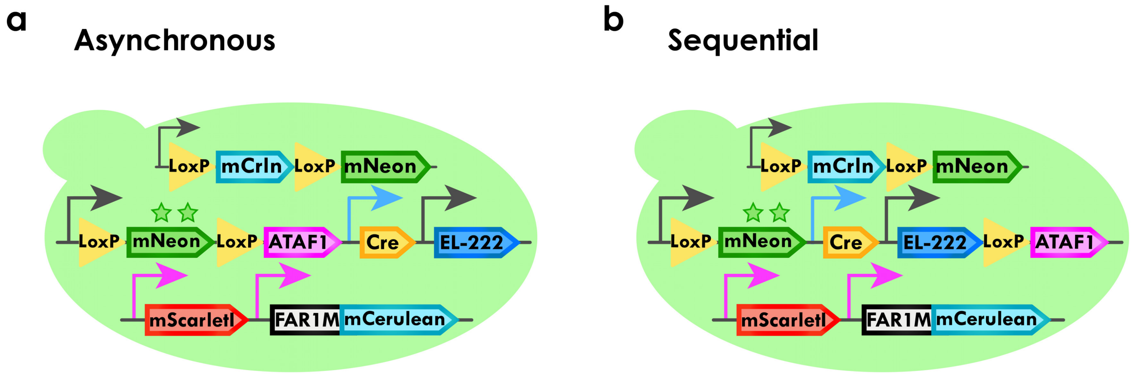


*Figure S7.2 Circuits for* ***(a)*** *asynchronous differentiation program and* ***(b)*** *sequential differentiation program which result in 4 and 3 species microbial consortia, respectively.*

All four species could be observed for the asynchronous differentiation program while only 3 could be observed with the sequential differentiation program (*Figure S7.3*).

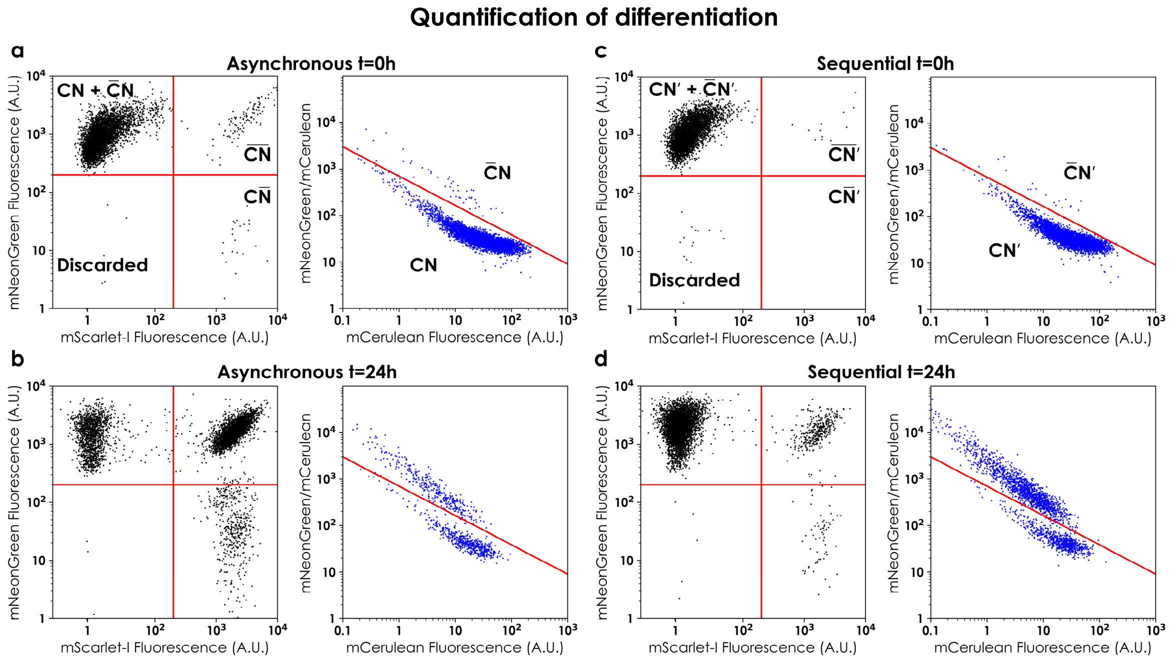


*Figure S7.3 Quantification of species prevalence. Cells were first separated by mScarlet-I fluorescence. Cells possessing red fluorescence were either* $\overline{CN}$ *or* $C\overline{N}$ *and those that did not were* $CN$ *or* $\bar{C}N$*. To distinguish between* $\overline{CN}$ *and* $C\overline{N}$ *mNeonGreen fluorescence was enough. However, to distinguish between* $CN$ *and* $\bar{C}N$*, both mNeonGreen and mCerulean fluorescence had to be analysed to obtain two well-separated populations (high mNeonGreen to mCerulean ratio corresponds to* $CN$ *and the lower to* $\bar{C}N$*). Cells not possessing appreciable levels of either mNeonGreen or mScarletI were discarded from analysis (<1%). The figure shows the four subpopulations before induction at t=0 and after stimulation with 3 light pulses of 30 minutes 6h apart at t=24h for both differentiation programs (Figure 5b & 5d).*

#### VIII. Note on handling of light sensitive strains

To obtain pure non-differentiated fraction, all cell manipulation was done in red light including transformation and the creation of stocks. Stocks were covered with aluminium foil during storage. Revivals were performed in red light and yeast plates were encased in aluminium foil during growth in the incubator. Overnight cultures were grown in the dark using a falcon holder covered on all sides. Overnight cultures were measured using cytometry prior to the start of preculture to ensure maximum non-differentiated cells. Typically, we observed 0.03%-1% differentiation at this point (for GAuDi we always observed 10%-20% background differentiation but as noted in the Section IV, we believe these cells were present due to leaky expression of ATAF1 TF and not background recombination). Some picked colonies were completely recombined and not used for experiments. We observed that streaked plates more than a week old led to higher levels of background differentiation in individual colonies. We note that the colonies were extremely sensitive to extraneous light and even opening them in red light led to significant increase in differentiated fraction over the next 96h (5%-10%). Such susceptibility to extraneous light was not observed for liquid cultures. Consequently, only freshly streaked plates were used for the experiments.
